# Fate of a supergene in the shift from diploidy to polyploidy

**DOI:** 10.1101/2025.01.18.633704

**Authors:** Emiliano Mora-Carrera, Narjes Yousefi, Giacomo Potente, Rebecca Lynn Stubbs, Barbara Keller, Étienne Léveillé-Bourret, Grob Stefan, Ferhat Celep, Giorgi Tedoradze, Elena Conti

## Abstract

Despite the evolutionary importance of supergenes, their properties in polyploids remain unexplored. Polyploid genomes are expected to undergo chromosomal rearrangements and gene losses over time, potentially affecting supergene architecture. The iconic distyly supergene (*S*-locus), controlling a floral heteromorphism with two self-incompatible morphs, has been well-documented in diploids, but remains unknown in polyploids. *Primula*, the classic model for distyly since Darwin, is ancestrally diploid and distylous, yet polyploid, homostylous species with a single, self-compatible floral morph evolved repeatedly. The intraspecific loss of distyly is associated with small loss-of-function mutations in the *S*-locus *CYP^T^* gene controlling style length and female self-incompatibility. Over longer timescales, relaxed selection on *CYP^T^* should generate greater accumulation of larger mutations, including exon and gene loss. By analyzing the first assembled genome of an allotetraploid, homostylous species (*Primula grandis*) in a comparative framework, we discovered two, nearly identical *S*-locus alleles in the same subgenome, suggesting it originated via inter-specific hybridization between a homostylous and a distylous progenitor. Conformant to predictions from theory, the macroevolutionary loss of distyly coincided with considerable degeneration of *CYP^T^*, while other *S*-locus genes remained largely unaffected, suggesting the shift to homostyly preceded and facilitated polyploid establishment. At the whole-genome level, we found minimal subgenome dominance — as expected, given the inferred recent origin of *P. grandis* — and highly reduced genetic diversity, congruently with its narrow distribution and self-compatibility. This study provides the first comparison of a supergene across ploidy levels and reproductive systems, contributing new knowledge on the previously unknown fate of supergenes in polyploids.

**SIGNIFICANCE:** This study advances knowledge on genome evolution by elucidating how supergenes (clusters of tightly linked genes) evolve across species with different sets of chromosomes and reproductive systems. By analyzing the newly assembled genome of the polyploid, self-compatible *Primula* grandis in a broad framework, we provide the first comparison of the distyly supergene between diploid outcrossers and polyploid self-fertilizers. We discovered one pair of identical supergene alleles in the same subgenome, rather than one pair per subgenome, revealing the species originated via a cross between a self-compatible and a self-incompatible progenitor. Conformant to theory, the gene controlling female self-incompatibility and style length (*CYP^T^*) was considerably degenerated, because of relaxed selection over time, with the rest of the supergene largely unaffected.

## INTRODUCTION

Transitions from diploidy to polyploidy and from outcrossing to selfing frequently co-occur in plants, attracting considerable attention from researchers across evolutionary biology and agronomy (Soltis & Soltis, 1999, 2000; Wendel & Cronn, 2003; Adams & Wendel, 2005; Igic *et al*., 2006; Flagel & Wendel, 2009; Robertson *et al*., 2011; Jiao *et al*., 2011; Paterson *et al*., 2012; van de Peer *et al*., 2017; Nieto Feliner *et al*., 2020; Yuan & Song, 2023). Selfing facilitates reproduction when mates are scarce, favoring the establishment of new polyploids and likely explaining why polyploids are often self-compatible (Otto & Whitton, 2000; Barringer, 2007; Bachmann *et al*., 2021; Novikova *et al*., 2023). *Primula* L. (primroses) is an ideal system to study the above-mentioned transitions because it includes both diploid species that are typically self-incompatible outcrossers and polyploid species that can self-fertilize (Richards, 2003).

Primroses have served as a classic evolutionary model since Darwin’s (Darwin, 1877) seminal studies on heterostyly, a floral heteromorphism characterized in *Primula* by two self-incompatible floral morphs: short-styled flowers (thrums) with short style and high anthers and long-styled flowers (pins) with long style and low anthers (i.e., distyly; **Fig. 1a**). An adaptation for efficient cross-pollination and outcrossing, heterostyly evolved repeatedly in angiosperms (Lloyd & Webb, 1992; Barrett, 2002; Naiki, 2012; Keller *et al*., 2014; Armbruster *et al*., 2017; Simón-Porcar *et al*., 2024). This trait increases genetic variation and species resilience (de Vos *et al*., 2014a) but requires mates and pollinators for reproduction. Conversely, homostyly — characterized by self-compatible, monomorphic flowers with anthers and stigma at the same level — enables selfing, thus decreasing genetic variation but affording reproduction when mates or pollinators are scarce (de Vos *et al*., 2012, 2014b, 2018), for instance after long-distance dispersal or range expansion into uncolonized habitats (Baker, 1955; Husband & Barrett, 1991). In *Primula*, homostyles typically have long style and high anthers (**Fig. 1a**). Phylogenetic analyses inferred the ancestor of the *Primula* clade (500+ spp.) to be distyslous and diploid (de Vos *et al*., 2014a), while homostylous, likely allopolyploid taxa evolved repeatedly via secondary contact following Pleistocene glacial retreat (Guggisberg *et al*., 2006, 2008, 2009). Conversely, the autopolyploid origin of homostyles has rarely been proposed (Dowrick, 1956).

**Fig. 1.**
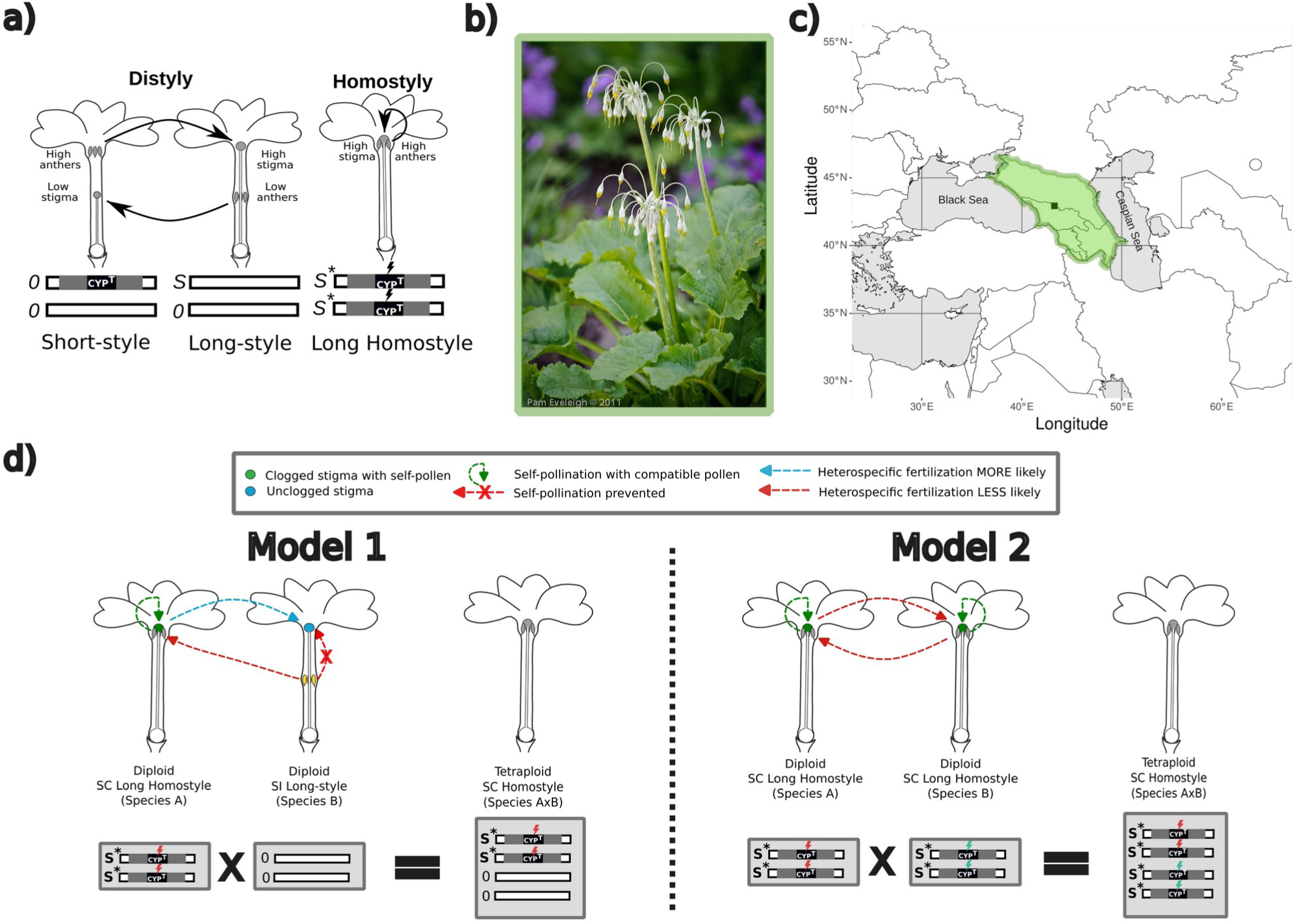
Shift from diploid distyly to polyploid homostyly in *Primula*. **a)** Distylous and homostylous phenotypes and *S*-locus genotypes in diploid *P. vulgaris*. Top, from left to right: Short-styled (thrum) and long-styled (pin) flowers: reciprocal positioning of male and female sexual organs in the two floral morphs promotes cross-pollination (solid arrows), while self-incompatibility (SI) prevents selfing, thus heterostyly favors outcrossing. In *Primula* sect. *Primula*, the heterostyly *S*-locus is hemizygous and present exclusively in short-styled (*S*/0) individuals, with *CYP^T^*determining short styles and female incompatibility; the *S*-locus is absent from long-styled (*0*/*0*) individuals. Bottom: Long-homostylous flower with high stigma and anthers in close proximity, favoring self-pollination; long-homostyles typically have two copies of the *S*-locus with a disrupted *CYP^T^* (*S\**/*S\**), enabling self-fertilization (Mora-Carrera et al., 2023); **b)** Inflorescence and single flower of polyploid, homostylous *P. grandis* (pictures courtesy of Pam Eveleigh © 2011); **c)** Distribution map of *P. grandis*, endemic (dark green square) to the Caucasus mountain range (outlined in lighter green): two populations were sampled; **d)** Models for the expected number of S-loci in the presumably allotetraploid *P. grandis:* under Model-1 (inter-specific cross between a homostyle and a long-styled plant) only two alleles of the *S*-locus are expected in the same subgenome; under Model-2 (inter-specific cross between two homostyles) four alleles of the *S*-locus are expected, two per subgenome, with different (marked in red and blue for Species A and B, respectively) mutations between subgenomes; Model-1 is hypothesized to be more likely due to less stigma clogging by self-pollen in the long-styled flower. Arrows indicate the direction of pollination from anthers to stigma.

Distyly is typically controlled by a hemizygous supergene present only in thrums but absent from pins (Li *et al*., 2016; Gutiérrez-Valencia *et al*., 2022; Raimondeau *et al*., 2024). Supergenes are groups of tightly linked genes inherited as a single unit due to suppressed recombination; they control multi-trait, balanced polymorphisms that are central to key evolutionary processes such as adaptation and reproduction (Schwander *et al*., 2014; Berdan *et al*., 2022). The heterostyly (or distyly) supergene, also known as the *S*-locus, has been most thoroughly studied in *Primula*, where it comprises 4-5 genes called *S*-genes hereafter (Li *et al*., 2016; Huu *et al*., 2020; Potente *et al*., 2022a, 2024). Function has been experimentally characterized for only two of these genes: *CYP^T^* determines short style and female incompatibility, while *GLO^T^* determines high anthers in thrums of distylous primroses (**Fig. 1a**; Huu *et al*., 2016, 2020). Within the ancestrally distylous, diploid *Primula vulgaris* Huds., homostylous, self-compatible plants evolved repeatedly from self-incompatible thrums via different loss-of-function mutations in *CYP^T^* (Mora-Carrera *et al*., 2023). *Primula vulgaris* homostyles co-occurring with pins and thrums have either nonsynonymous mutations or small deletions in *CYP^T^*; a large deletion comprising exon-1 was found only in homostyles from the single known monomorphic population. Due to self-fertilization, most homostyles of *P. vulgaris* have two, rather than one, disrupted *CYP^T^* copies (Mora-Carrera *et al*., 2024).

Despite the above-mentioned infraspecific studies, the fate of *CYP^T^*and the heterostyly supergene in interspecific shifts to polyploid homostyly remains unknown. Theory predicts that, over macroevolutionary timescales, relaxed selection on genes associated with lost traits leads to the accumulation of extensive mutations, including large insertions and/or deletions, exon loss, or entire gene loss (Collin & Miglietta, 2008; Lahti *et al*., 2009). Once a gene loses its function, it becomes a pseudogene, a nonfunctional remnant of a gene that was once active (Cheetham *et al*., 2020). The fate of pseudogenes over time, including their potential loss, depends on environmental pressures, selection, pleiotropic effects, and the possibility of being repurposed for similar or other functions (Cheetham *et al*., 2020). Does extensive degradation and even loss of *CYP^T^* occur in interspecific shifts to homostyly, as predicted? Furthermore, theory predicts that polyploid genomes rediploidize over time, a process involving extensive chromosomal rearrangements and gene losses that could also affect the *S*-locus (Li *et al*., 2021). Does the transition from diploidy to polyploidy affect the organization and copy number of the heterostyly supergene? Overall, the impact of polyploidy on supergenes remains unknown.

To fill these gaps of knowledge, we focus on the tetraploid *Primula grandis* Trautv. (2n=4x=22) because its biological characteristics and available resources are ideal to clarify the macroevolutionary consequences of shifts to polyploidy on supergenes (**Fig. 1b,c**). *Primula grandis*, narrowly endemic in the Caucasus region, is the only polyploid (Brunn, 1932), homostylous, self-compatible species of *Primula* sect. *Primula*, a small, young clade (ca. 2.46 million years old; de Vos *et al*., 2014a) additionally comprising six diploid, heterostylous species all sharing the same base chromosome number (x=11; Richards, 2003). The conflicting placement of *P. grandis* in phylogenies inferred from few nuclear and chloroplast DNA loci (Schmidt-Lebuhn *et al*., 2012) and Whole Genome Sequencing (WGS) data (Stubbs *et al*., 2023) suggests an allotetraploid origin, but this hypothesis has never been tested via the assembly and analysis of its entire genome. Genome assemblies are available for two diploid, heterostylous relatives of *P. grandis*: a high-resolution, chromosome-scale assembly for *Primula veris* L. (Potente *et al*., 2022a) and a fragmented draft assembly for *P. vulgaris* (Cocker *et al*., 2018). Furthermore, extensive WGS data from all species of *Primula* sect. *Primula* are available (Stubbs *et al*., 2023, 2024).

By integrating analyses of the here assembled, annotated genome of *P. grandis* with newly and previously generated WGS data in a broad comparative framework, we provide the first study of the fate of the hemizygous heterostyly supergene in a polyploid genomic background. We test the two main models for the expected number of *S*-loci in a homostylous, presumed allotetraploid species (**Fig. 1d**). If the species stemmed from a cross between a homostyle and a long-styled plant, two nearly homozygous alleles of the *S*-locus with the same disruptive mutations in *CYP^T^*are expected in the same subgenome (Model-1). If the species stemmed from a cross between two homostyles, four copies of the *S*-locus — two nearly homozygous alleles per subgenome with likely different disruptive *CYP^T^* mutations between subgenomes — are expected (Model-2). Importantly, homostyles are cross-compatible with long-styled but not short-styled plants, limiting the possibilities for the initial cross generating a polyploid, homostylous species (Lewis & Jones, 1992). We also test the prediction that the interspecific loss of heterostyly is associated with degradation and possibly loss of *CYP^T^*but has little or no effect on the remaining *S*-genes. Furthermore, given the young age of the *Primula* sect. *Primula* clade and the phylogenetically proposed allopolyploid origin of *P. grandis* (Schmidt-Lebuhn *et al*., 2012; Stubbs *et al*., 2023), we hypothesize its genome to comprise two distinct subgenomes scarcely affected by rediploidization and subgenome dominance, as predicted by theory (Wendel, 2015; Feng *et al*., 2024). We then test whether any extant species served as progenitors of *P. grandis* by inferring genome-wide phylogenies of *Primula* sect. *Primula*. Additionally, theory predicts that polyploid speciation should cause strong genetic bottlenecks (Stebbins, 1950) leading to sharp decrease of genome-wide genetic diversity and reduced efficacy of purifying selection (Hough *et al*., 2013). Finally, genetic redundancy associated with polyploidization should relax selection on duplicated genes (Ohno, 1970). We test the above predictions on *P. grandis* via demographic and population genomic analyses across *Primula* sect. *Primula*. The here-presented first genome assembly of a tetraploid, homostylous species, analyzed within a comparative genomic framework, contributes new knowledge on the genomics of shifts between ploidy levels and breeding systems and the previously unknown fate of supergenes in polyploid genomes.

## RESULTS

### Chromosome-scale assembly of *P. grandis* tetraploid genome

The size of the *P. grandis* tetraploid genome estimated via flow-cytometry (898 Mb) was about twice (x 1.9) that of the closely related diploid *P. veris* and *P. vulgaris* (452 and 430 Mb, respectively; Cocker *et al*., 2018; Potente *et al*., 2022a). To assemble a high-quality reference genome for *P. grandis*, long- and short-read sequences were combined with Hi-C data for scaffolding (**Figs. S1, S2, S3**). The size of the assembled genome (768.24 Mb; N50 = 32.92 Mb; **Fig. 2a**) fell between the sizes estimated via *k*-mer analysis of short-reads (697.70 Mb) and flow cytometry (∼898 Mb; **Fig. S4**). The high repetitive content (∼52%; **Fig. S8**) of the *P. grandis* genome might explain the bioinformatic underestimation of genome size. The BUSCO completeness score (94.4%) supported the completeness of the genome assembly (**Fig. S5**). The final assembly produced 2134 scaffolds, of which 22 encompassed 743.54 Mb (96.78%) of the total assembly size, corresponding to the 22 chromosomes of *P. grandis* (**Fig. 2a, b**). The gene annotation identified 40,882 protein-coding genes distributed along the 22 chromosome-scale scaffolds (**Table S2**).

**Fig. 2.**
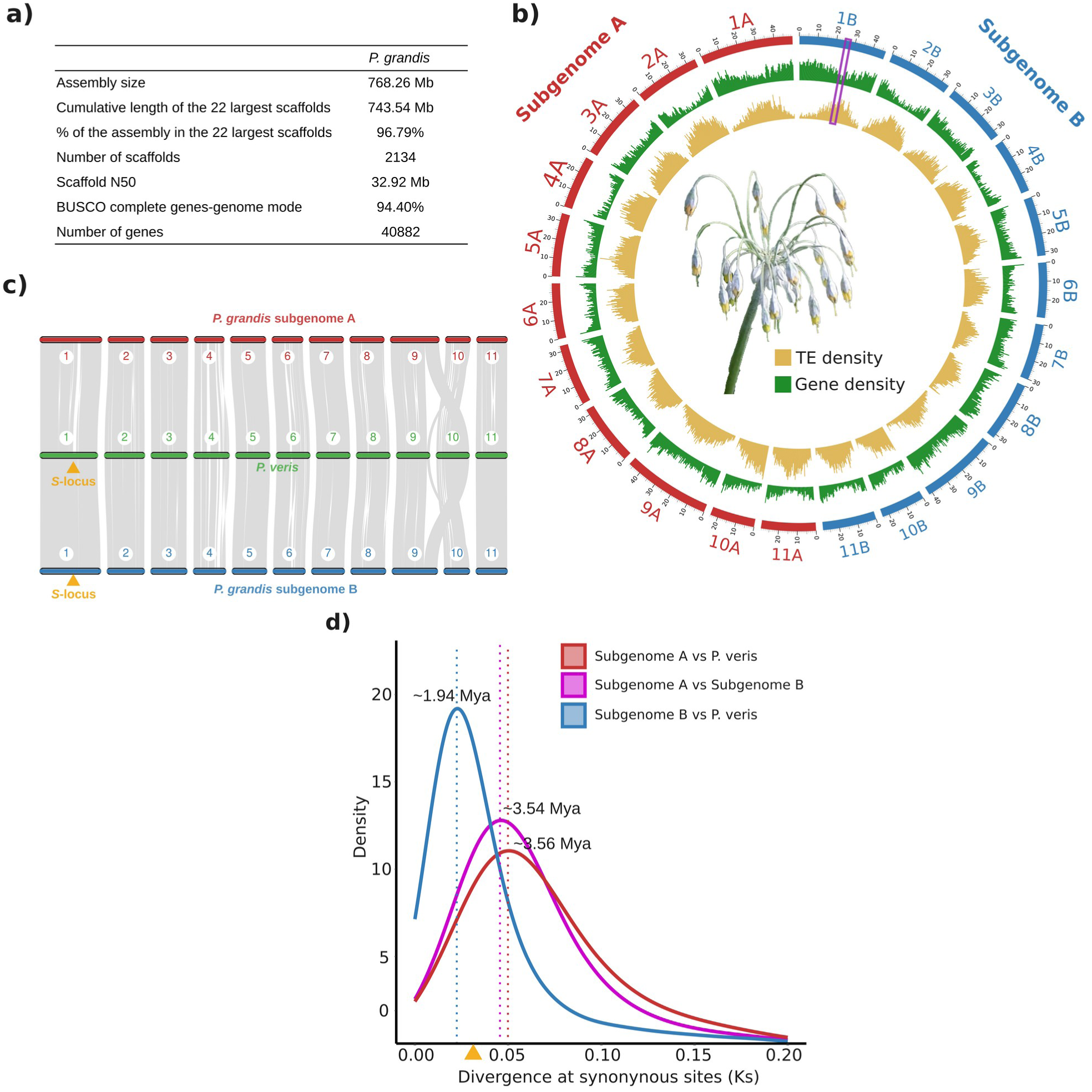
Genome assembly and subgenome characterization of the tetraploid, homostylous *Primula grandis*. **a)** Summary statistics of the *P. grandis* genome assembly and gene annotation. **b)** Circos plot of the *P. grandis* chromosome-scale genome assembly showing the 22 largest scaffolds assigned to subgenome A and B, in red and blue, respectively; distributions of Transposable Elements (TE) and gene density along each chromosome are represented in yellow and green, respectively; the purple rectangle in chromosome 1B indicates the position of the *S*-locus. **c)** Collinearity analyses between the 11 chromosomes each in subgenomes A and B of *P. grandis* and the 11 chromosomes of *P. veris*: the *S*-locus (marked by a yellow triangle) is found exclusively in subgenome B of *P. grandis* and in the same chromosome (ch. 1) in both *P. grandis* and *P. veris*. **d)** Density distributions of divergence at synonymous sites (*Ks*) of paralogous gene copies between *P. veris* and both subgenomes of *P. grandis*. Dotted lines indicate peaks in the distribution of *Ks* with estimated divergence times between species in Million years ago (Mya). Crown age of *Primula* sect. *Primula* (∼2.46 million years ago; de Vos et al., 2014) is marked by a yellow triangle.

**Fig. 3.**
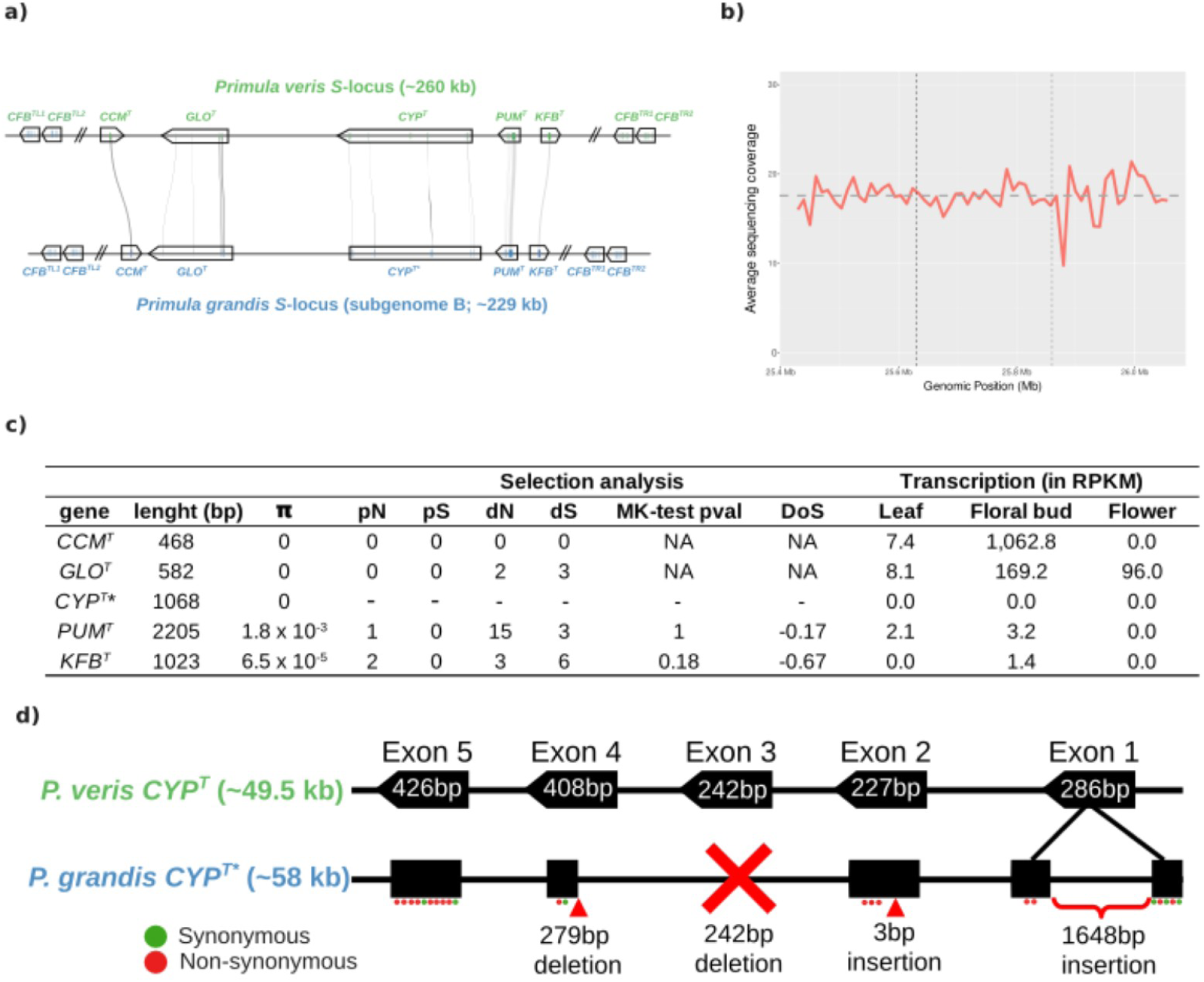
Characteristics of the *S*-locus supergene in a polyploid genome. **a)** Microsynteny analysis showing high collinearity between *Primula veris* and *P. grandis S*-loci: the grey lines between *S*-loci connect the exons of each of the fives S-genes (*CCM^T^*, *GLO^T^*, *CYP^T^*, *PUM^T^*, and *KFB^T^*); the length of the *S*-locus is 276kbp in *P. veris* and 229 kbp in *P. grandis*; in both species the *S*-locus is flanked by two copies of the *CFB* gene on each side. **b)** Coverage analysis showing no changes across the *S*-locus and its up- and down-stream 200 kbp flanking regions in subgenome-B of *P. grandis*; plot shows 10 kb windowed average sequencing coverage of the *S*-locus (y-axis) based on Whole Genome Resequencing (WGR) data from 10 *P. grandi*s individuals; horizontal grey dashed line indicates average sequencing depth across chromosome-1 of *P. grandis* subgenome-B (average = 17.59); vertical black dashed lines indicate genomic limits of *S*-locus. **c)** Selection and transcription analyses in *P. grandis S*-genes: Length (in basepairs, bp) per gene, estimates of nucleotide diversity (π), number of non-synonymous polymorphic sites (pN), number of synonymous polymorphic sites (pS), number of non-synonymous fixed sites (dN), number of synonymous fixed sites (dS), significance values of McDonald-Kreitman tests (MK-test pval), and Direction of Selection (DoS, with negative values indicating accumulation of slightly deleterious mutations); Expression levels in leaf, floral bud, and flower tissues of *P. grandis* expressed as Reads Per Kilobase per Million (RPKM); *indicates pseudogene; NA=Not Applicable. **d)** Structure of functional *CYP^T^ S*-gene (1587 bp long, five exons with bp length in white) in *P. veris* and disrupted *CYP^T^*pseudogene (1068 bp recovered, exon 3 missing, insertions and deletions in exons 1, 2, 4) in *P. grandis*. Green and red circles indicate synonymous and non-synonymous mutations in each exon when compared to *P. veris CYP^T^*. Exon and intron sizes are not to scale.

### Subgenomes characteristics

The *k*-mer-specific analysis of repetitive elements enabled the assignment of the 22 chromosome-scale scaffolds to two clearly distinct subgenomes (subgenomes-A and -B, hereafter; **Fig. S6**), each containing 11 chromosomes (**Fig. 2b**) that were highly collinear with the 11 chromosomes of *P. veris* (**Figs. 2c, S7, S8**). Subgenomes-A and -B comprised 380.64 Mb and 362.90 Mb, respectively, and the 17.73 Mb size-difference between subgenomes was evenly distributed along the 11 homoeologous chromosomes (**Table S2**). Annotation of repetitive elements revealed that subgenome-A comprised a slightly higher proportion of TEs than subgenome-B (45.63% vs. 40.95%, respectively; **Fig. S9**). Furthermore, subgenome-A had a lower number of genes (19,976 *vs*. 20,906, respectively; **Table S2**) and lower gene density than subgenome-B (**Fig. S10**). Transcriptome analyses indicated minor homoeolog-expression bias (HEB) towards subgenome-B in leaves (n_sub_a_=245 vs n_sub_b_ 278 gene pairs), floral buds (n_sub_a_ 235 vs n_sub_b_ 258 gene pairs), and flowers (n_sub_a_ 276 vs n_sub_b_ 316 gene pairs) (**Fig. S11**). Estimates of genome-wide divergence at synonymous (syn) sites (*Ks*) between subgenomes-A and -B showed a peak at *Ks*=0.04, corresponding to a divergence time of ∼3.56 Mya (CI=3.91-3.31 Mya; **Fig. 2d**). Furthermore, *Ks* values of *P. veris* vs. subgenomes-A and -B, respectively, were 0.04 and 0.02, corresponding to divergence times of ∼3.54 Mya (CI=3.88-3.28 Mya) between subgenome-A and *P. veris* and ∼1.94 Mya (CI=2.13-1.80 Mya) between subgenome-B and *P. veris* (**Fig. 2d)**.

### Characteristics of the S-locus in the tetraploid *P. grandis*

To identify the *P. grandis S*-locus, we used BLAST to screen the *P. grandis* genome for the five *S*-genes and the four *CFB* genes flanking the *S*-locus (*CFB^TL1^* and *CFB^TL2^*on the left and *CFB^TR1^* and *CFB^TR2^* on the right) in *P. veris* (**Fig. 3a**). These nine genes were found in a ∼229-kb region — characterized by a high-density of TEs — of chromosome 1 in subgenome-B (**Fig. 2b**) in the same order and orientation as in *P. veris* (**Fig. 3a**). Synteny analyses showed that collinearity between *P. veris* and *P. grandis* extended well beyond the *S*-locus in chromosome 1 (**Fig. 2c**). Coverage analysis of the *P. grandis* genome using WGS data indicated that the *S*-locus is present in both haplotypes (i.e., non-hemizygous) of chromosome 1B (**Fig. 3b).**

Nucleotide diversity in *P. grandis S*-genes was zero in *CCM^T^*, *GLO^T^* and *CYP^T^* and very low in *PUM^T^* (1 nonsynonymous and 0 synonymous polymorphic sites) and *KFB^T^* (2 nonsynonymous and 0 synonymous polymorphic sites; **Fig. 3c**), indicating low *S*-locus differentiation in the young, self-compatible, narrow Caucasus endemic *P. grandis*. Sequence comparisons between the *CCM^T^*, *GLO^T^*, *PUM^T^*, and *KFB^T^ S*-genes of the homostylous *P. grandis* and those of the heterostylous *P. veris* detected no loss-of-function mutations for any of these genes in the former species. Indeed, the four genes were all transcribed, hence presumably functional, in *P. grandis* floral buds (**Fig. 3c**). The highest accumulation of nonsynonymous mutations (15) among the four genes above occurred in *PUM^T^*, compared to only 3 synonymous mutations (**Fig. 3c**). Furthermore, McDonald-Kreitman (MK) tests comparing *S*-genes between *P. grandis* and *P. veris* (which could not be performed on *CCM^T^*, *GLO^T^* and *CYP^T^* due to absence of polymorphic and/or fixed sites) did not detect significantly reduced efficacy of purifying selection on *P. grandis PUM^T^* and *KFB^T^* (**Fig. 3c**). Direction of Selection (DoS) analyses, also applicable only to *PUM^T^* and *KFB^T^*, revealed low accumulation of slightly deleterious mutations in these two genes. Finally, we found evidence of extensive degeneration in *P. grandis CYP^T^* via the accumulation of the following segmental and point mutations compared to the functional *P. veris* gene(**Fig. 3d**): two insertions (a 1648bp insertion in exon-1 and a 3bp insertion in exon-2); two large deletions (loss of exon-3 and loss of 279 bps in exon-4); and 16 nonsynonymous mutations (four in exon-1, three in exon-2, one in exon-4, and eight in exon-5; **Table S3**). Congruently, *CYP^T^* was not expressed in leaf, floral-bud or flower transcriptomes (**Fig. 3d**), confirming *CYP^T^* pseudogenization in *P. grandis*. No variation was found in *CYP^T^* pseudogene of 10 *P. grandis* individuals. The total length of *P. grandis CYP^T^* pseudogene was ca 58 Kb, the cumulative length of its exons was 1068 bp (**Fig. 3d**).

### Concurrent bottlenecks in both *P. grandis* subgenomes

The results of the stairway plots indicated that both *P. grandis* subgenomes had reduced effective population size, implying recent bottlenecks (**Fig. S12**). Furthermore, model-based coalescent demographic analyses with *fastsimcoal2* supported overlapping genetic bottlenecks in subgenomes-A and -B (**Fig. S13)** (T_BOT_=25,054 [95% CI: 20,726-29,382] and T_BOT_=20021 [95% CI:15,830-24,211] generations ago, respectively; **Fig. 4a**), implying that the two subgenomes came together via allopolyploid speciation 31,660-58,764 ybp, assuming 2 years generation time for *P. grandis* (**Fig. 4a**). The bottlenecks were confirmed by more positive Tajima’s D values for both *P. grandis* subgenomes compared to other diploid species of *Primula* sect. *Primula* sampled from the Caucasus, except for *P. renifolia*, suggesting a bottleneck also in this species (**Fig. 4b**).

**Fig. 4.**
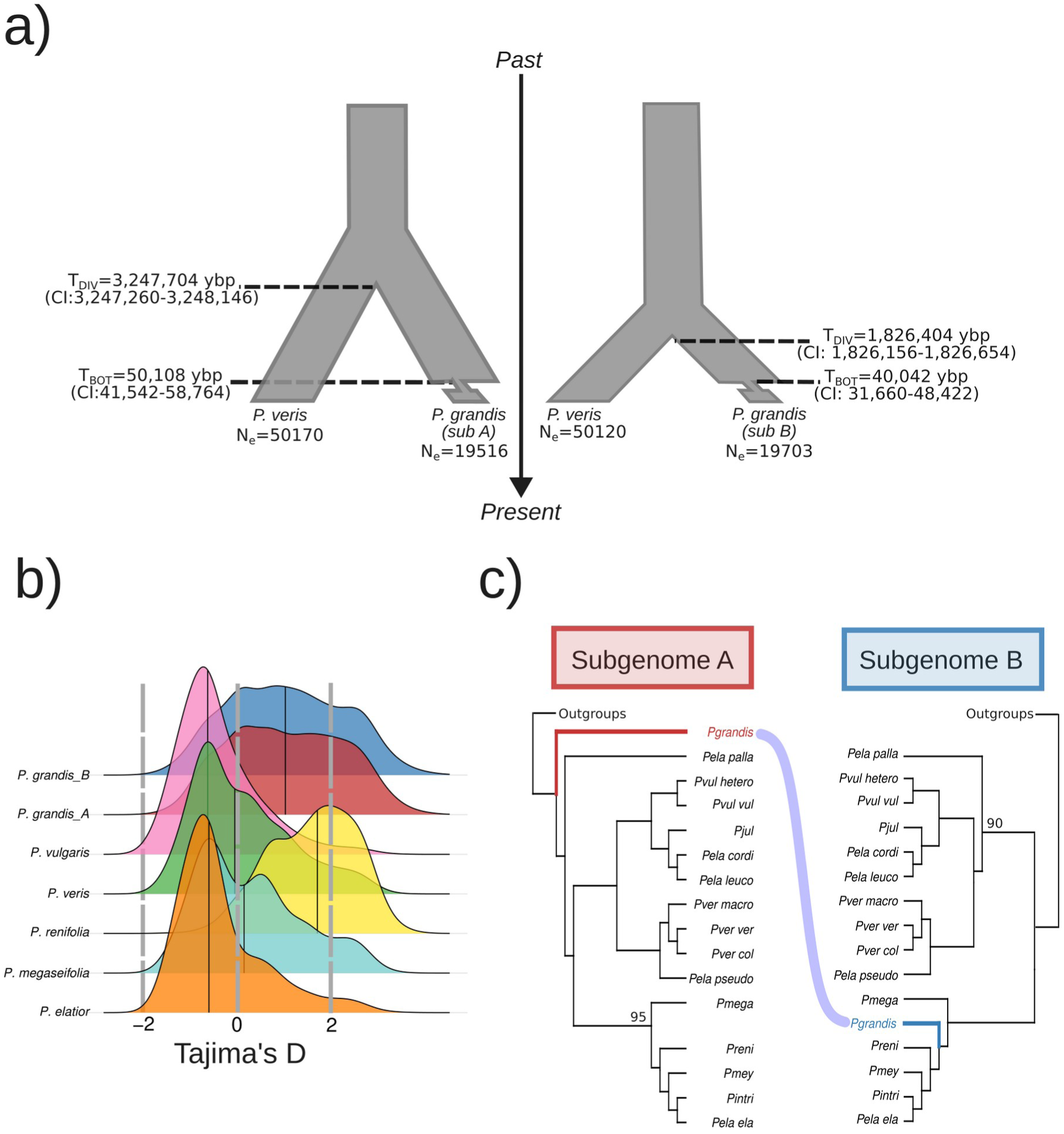
Origin of *Primula grandis* and its subgenomes. **a)** Best-supported demographic models inferred with coalescent simulations (see Materials and Methods; Fig. S13) showing the estimated times (with 95% Confidence Intervals) of divergence between *P. veris* and subgenomes-A (left) and -B (right) of *P. grandis* (T_DIV_) reported in years before present (ybp), assuming two-year generation time for *P. grandis*, and of the genetic bottleneck in each subgenome (T_BOT_); *N_e_* indicates effective population size. **b)** Distribution of Tajima’s D estimates for *P. grandis* subgenomes-A and -B (red and blue, respectively), *P. vulgaris* (pink), *P. veris* (green), *P. renifolia* (yellow), *P. megaseifolia* (turquoise), and *P. elatior* (orange). Black lines within distributions denote median values of Tajima’s D; positive Tajima’s D values indicate genetic contraction, negative values indicate population expansion. **c)** Tanglegram of genome-wide Maximum Likelihood (ML) phylogenies inferred from collinear orthologues showing the relationships of *P. grandis* subgenomes-A and -B (red and blue, respectively) within *Primula* sect. *Primula*. Species codes are as follows: *P. elatior* (*Pela_cordi*=*P. elatior_*subsp. *cordifolia, Pela_ela=P. elatior* subsp. *elatior, Pela_ela=P. elatior* subsp. *leucophylla, Pela_palla=P. elatior* subsp. *pallasi,* and *Pela_pseudo=P. elatior* subsp. *pseudoelatior*); *Pintri=P. intricata; Pmey=P. meyeri; P. veris* (*Pver_col=P. veris* subsp. *columnae, Pver_macro=P. veris* subsp. *macrocalyx*, *Pver_ver=P. veris* subsp. *veris*); *Primula vulgaris* (*Pvul_hetero=P. vulgaris* subsp. *heterochroma* and *Pvul_vul*=*P. vulgaris* subsp. *vulgaris*); *Pgrandis=P. grandis, Pmega=P. megaseifolia, Pjul=P. juliae* and *Preni=P. renifolia*. Outgroups outside *Primula* sect. *Primula* are not shown. Nodal bootstrap support (BS) values lower than 100% are reported; all other nodes received 100% BS. See Fig. S14 for complete phylogenies.

### Origin of the allopolyploid *P. grandis*

Phylogenetic analyses revealed that subgenomes-A and -B from 10 individuals of *P. grandis* formed two strongly supported monophyletic groups (**Fig. S14)**. Subgenome-A was strongly supported as sister to the remaining members of the *Primula* sect. *Primula* clade and subgenome-B was weakly supported as sister to the clade comprising *Primula renifolia* Volgunov, *Primula elatior* subsp. *elatior*, *Primula intricata*, and *Primula meyeri* (**Figs. 4C, S15**), suggesting a more recent origin of P. grandis subgenome-B than subgenome-A, consistently with the divergence times estimated above **(Fig. 2d**).

### Population genomic consequences of polyploidization in *P. grandis*

*Primula grandis* had the lowest genome-wide diversity, as estimated by S_syn_, and π_syn_, of the *Primula* sect. *Primula* species in the Caucasus, except for *P. renifolia* (**Fig. 5a**). Within *P. grandis*, both subgenomes showed similar values of nucleotide diversity (mean π_syn_ ± sd; 0.0008 ± 0.001 and 0.0008 ± 0.001 for subgenomes-A and -B, respectively; **Fig. 5a**). Furthermore, LD decay in *P. grandis* was slower than in *P. vulgaris, P. veris*, and *P. megaseifolia*, but faster than in *P. renifolia* and *P. elatior* from Caucasus (**Fig. S15**). The volcano plots based on MK-tests indicate that both subgenomes of *P. grandis* had significantly more genes with negative values of Direction of Selection (DoS) than the genomes of diploid relatives (**Fig. 5b**). Within *P. grandis*, subgenome-A had eight more genes with significantly less efficient purifying selection (i.e., DoS < 0) and five more genes evolving under positive selection (i.e., DoS > 0) than subgenome-B. Finally, when one gene was under relaxed selection in one subgenome, its homoeologous copy in the other subgenome appeared to be constrained by selection (**Table S4**).

**Fig. 5.**
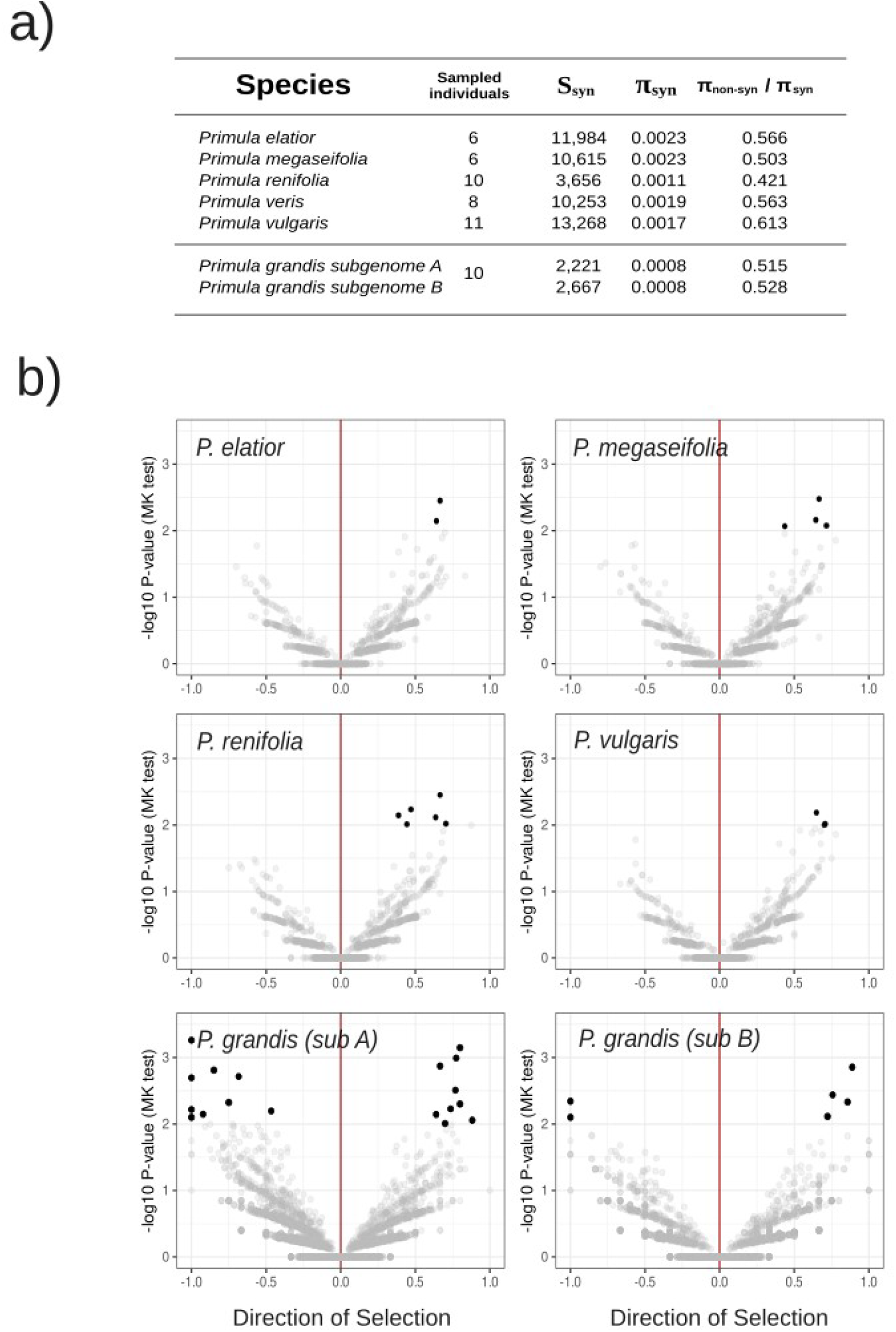
Genetic diversity and selection in the tetraploid, homostylous *Primula grandis*. **a)** Summary of individuals per species of *Primula* sect. *Primula* sampled in the Caucasus and genome-wide diversity estimates including number of segregating sites (S_syn_), pairwise nucleotide diversity (π_syn_) at synonymous (4-fold degenerate) sites, and the ratio π of non-synonymous (non-syn; 0-fold degenerate) to synonymous sites. All values, except S_syn_, represent the genome-wide average over 50,000 non-overlapping windows; **b)** Volcano plots based on MK-tests depicting the direction of selection (DoS) in four diploid species of *Primula* sect. *Primula* and the tetraploid *P. grandis* [subgenome-A and subgenome-B]; negative DoS values indicate accumulation of slightly deleterious mutations, positive values indicate accumulation of slightly beneficial mutations; DoS values of zero (red line) indicate neutral mutations. Gene transcripts with significant (log_10_*P* = 2 in the Y-axis; *P* <0.01) deviation from neutrality based on MK-tests are marked by black dots; grey dots indicate non-significant deviation from neutrality in MK-tests. *Primula veris* was used as outgroup for the MK-test (see Material and Methods).

## DISCUSSION

Despite the evolutionary importance of supergenes, their characteristics in polyploid genomes remain unknown. Over time, polyploid genomes are expected to undergo extensive chromosomal rearrangements and gene losses (Li *et al*., 2021), raising questions about supergene organization and copy number in polyploids. By focusing on the heterostyly supergene, the present study enabled the first comparison of a supergene across ploidy levels and reproductive systems. While well-characterized in diploid genomes of several heterostylous (Cocker *et al*., 2018; Shore *et al*., 2019; Potente *et al*., 2022a; Gutiérrez-Valencia *et al*., 2022; Fawcett *et al*., 2023; Raimondeau *et al*., 2024; Potente *et al*., 2024) and one homostylous species (Gutiérrez-Valencia, 2024), its structure in polyploid genomes was unknown. In the tetraploid, homostylous *P. grandis*, we found the same five *S*-genes (with *CYP^T^* pseudogenized) in the same order, orientation, and chromosome (chr.1) as in the diploid, heterostylous *P. veris* (**Fig. 3b**; Potente *et al*., 2022a) and *P. vulgaris* (Li *et al*., 2016), although the *P. grandis S*-locus was slightly smaller (229 Kb *vs* 278Kb and 260 Kb, respectively). Conversely, the distantly related, diploid, heterostylous *Primula edelbergii* has a larger *S*-locus (633 Kb), containing only four genes (with *CCM^T^* missing) in a different order and chromosome (chr.6; Potente *et al*., 2024). These findings reveal that the organization of the *S*-locus in *Primula* is conserved across closely related species regardless of ploidy level and reproductive strategy, while diverging across larger evolutionary timescales.

The number, divergence, and subgenomic location of *S*-locus copies offer insights about the original cross giving origin to allopolyploid, homostylous species. We discovered that the tetraploid *P. grandis* contains only two nearly homozygous *S*-locus alleles in subgenome-B (**Figs. 2, 3**), rather than two distinct pairs (one per subgenome) of *S*-locus alleles (Models 1 and 2 in **Fig. 1d**). The minimal divergence and occurrence of the two *S*-locus alleles within the same subgenome suggest that *P. grandis* originated from an interspecific cross between a homostylous pollen-donor and a long-styled pollen-recipient, rather than between two homostylous parents (Models 1 and 2, respectively, in **Fig. 1d**). Indeed, it is expected that the stigmas of long-styled flowers are more likely to be fertilized by heterospecific pollen from homostyles because they are less clogged by self-pollen (Model 1, **Fig. 1**; Barrett, 2002; Keller *et al*., 2014). Moreover, pollen of homostyles should retain the compatibility characteristics of short-styled flowers, making it cross-compatible with long-styled stigmas. Conversely, crosses between homostyles and short-styled plants should be incompatible, as both are expected to produce pollen with the same incompatibility type (Wedderburn & Richards, 1992). Indeed, experimental crosses between homostyles and long-styled plants were found to produce more seeds than crosses between homostyles of different *Primula* species (Wedderburn & Richards, 1992). Importantly, the progeny of an allopolyploid cross between a homostyle and a long-styled plant inherit disrupted *CYP^T^*, enabling reproduction via selfing of the newly formed polyploids and species establishment. Lastly, research on Brassicaceae demonstrates that the loss of homomorphic, multi-allelic self-incompatibility in one progenitor facilitates reproduction and the establishment of new allopolyploids (Novikova *et al*., 2023), which aligns with our findings supporting the hypothesis that *P. grandis* originated from a self-compatible homostyle and a long-styled plant.

Theoretical research on relaxed selection has mainly focused on the loss of phenotypic traits (Lahti *et al*., 2009), rather than on their controlling genes (but see Somel *et al*., 2013; Zheng *et al*., 2022). Heterostyly, an adaptation for outcrossing, is often lost when mates or pollinators are scarce (Barrett, 2019). At the microevolutionary scale, loss of heterostyly within *P. vulgaris* was associated with different loss-of-function mutations in *CYP^T^*, including nonsynonymous mutations, small deletions and insertions in homostyles (Mora-Carrera *et al*., 2023, 2024). At the macroevolutionary scale, the *S*-locus of the young homostylous *P. grandis* (estimated age: 31,660-58,764 years; **Fig. 4a**) retains the *CYP^T^* pseudogene, but with extensive mutations, including the loss of exon-3, more than half of exon-4, and a 1648-bp deletion in exon-1 (**Fig. 3d**). Thus, evidence supports the conclusion that, as predicted from theory, relaxed selection on *CYP^T^*pseudogene allows for the accumulation of increasingly larger structural mutations in interspecific losses of heterostyly. Not surprisingly, *P. grandis CYP^T^* also accumulated the largest number (24) of point-mutations of all *S*-genes compared to *P. veris* (**Fig. 3b, c**). Contrariwise, no or very few such mutations were found in *CCM^T^*, *GLO^T^*, and *KFB^T^* (**Fig. 3c**). The only other *S*-gene with high numbers of point mutations was *PUM^T^*, with 15 nonsynonymous vs three synonymous mutations compared to *P. veris* (**Fig. 3c**). This result aligns with previously reported high ratios of divergence at nonsynonymous to synonymous sites in *PUM^T^* of *P. veris* and *P. edelbergii* (Potente *et al*., 2022a, 2024) suggesting relaxed selection on this gene. The low expression levels in both leaves and young floral buds of *PUM^T^*in the homostylous *P. grandis* (**Fig. 3c**) and in the heterostylous *P. vulgaris*, *P. veris*, and *P. edelbergii* (Cocker *et al*., 2018; Potente *et al*., 2022b, 2024) call into question whether *PUM^T^*plays a role in controlling any heterostylous trait. Functional analyses of this gene are necessary to elucidate this question. Altogether, the evidence supports the conclusion that the macroevolutionary loss of heterostyly is associated with pseudogenization and degeneration of *CYP^T^*, as predicted, while the rest of the *S*-locus remains largely unaffected. As comparative genomic analyses of trait-controlling genes expand, further insights into the complex interplay among repeated trait loss, relaxed selection, pseudogenization, and the dynamics of gene loss versus rescue will continue to emerge (Collin & Miglietta, 2008; Schrader *et al*., 2021; Jones *et al*., 2023; Moran *et al*., 2023).

Elucidating the origin and evolution of polyploids has long been a major goal of evolutionary biology and agronomy, but progress was hindered by the difficulty of sequencing polyploid genomes. The recent advent of Hi-C technology has facilitated the assembly and phasing of polyploid genomes, especially in crop plants, but also in non-model organisms (Burton *et al*., 2013; The International Wheat Genome Sequencing Consortium (IWGSC) *et al*., 2014; Sun *et al*., 2022; Peng *et al*., 2022; Ma *et al*., 2023). Indeed, Hi-C scaffolding enabled the high-quality assembly of the 22 chromosomes of the tetraploid *P. grandis* (**Figs. 2, S2**). The clear assignment of the 22 chromosomes to two distinct subgenomes (**Figs. 2, S6**), high collinearity between them (**Figs. 2C, S8**), and recent origin (∼32,000-59,000 years ago) estimated for *P. grandis* (**Fig. 4a**) are consistent with the expectation of minimal genomic rearrangements in recently formed allopolyploids (Wendel *et al*., 2016; Mason & Wendel, 2020; Yuan & Song, 2023). Furthermore, the slightly higher number of genes (**Table S2**), higher gene density (**Fig. S10**), and minor expression bias of subgenome-B (**Fig. S11**) — estimated to be younger than subgenome-A (**Fig. 2d**) — point to marginal dominance of subgenome-B over subgenome-A. Altogether, the evidence suggests that *P. grandis* is a recent allopolyploid with two distinct subgenomes showing low levels of rediploidization and subgenome dominance. As more high-quality polyploid and diploid genome assemblies of related species become available, they will help clarify the interactions among species age, rediploidization, and subgenome dominance (Schiavinato *et al*., 2021; He *et al*., 2021; Feng *et al*., 2024; Ma *et al*., 2024).

Key questions on polyploidy include the identification of parental origins of allopolyploid, self-compatible species and patterns of polymorphism and selection on their genomes. Our genome-wide phylogenetic analyses placed *P. grandis* subgenome-A as sister to the rest of *Primula* sect. *Primula* (**Fig. 4c**), consistently with its older estimated age (**Figs. 2d, 4a**), and subgenome-B as sister to a shallower clade comprising the Caucasus endemic *P. renifolia* and Caucasus populations of *P. elatior*, consistently with its younger estimated age. However, no extant diploid species were identified as possible progenitors of *P. grandis* (**Fig. 4c**). These findings are consistent with the incongruent relationships of *P. grandis* in previous phylogenetic analyses of a few nuclear and chloroplast genes (Schmidt-Lebuhn *et al*., 2012) and genome-wide WGS data (Stubbs *et al*., 2023).

As expected from theory, we found that the allopolyploid speciation event that gave origin to the self-compatible *P. grandis* was associated with a shared, recent (∼32,000–59,000 years ago) genetic bottleneck affecting both its subgenomes (**Figs. 4a, b, S12**). Indeed, both subgenomes of *P. grandis* had the lowest genome-wide diversity of all *Primula* sect. *Primula* species, except for the narrow Caucasus endemic *P. renifolia* (**Figs. 4, 5**). Also as expected, MK-test analyses indicated that both subgenomes of *P. grandis* have reduced efficacy of purifying selection compared to heterostylous, diploid closely related species. Furthermore, the redundancy of polyploid genomes is thought to relax selection, allowing for the accumulation of slightly deleterious and beneficial mutations more readily than in diploids (Ohno, 1970; Comai, 2005; van de Peer *et al*., 2017; Paape *et al*., 2018). Conformingly, the subgenomes of *P. grandis* accumulated significantly more mutations of both types compared to closely related, diploid genomes (**Fig. 5b**). Genes under significantly less efficient purifying selection were found exclusively in the subgenomes of the tetraploid *P. grandis*, but not in the diploid genomes of *Primula* sect. *Primula* (**Fig. 5b**). Consistent with the younger age and inferred marginal dominance of subgenome-B, subgenome-A harbored both more deleterious and beneficial mutations, with only two genes under less efficient purifying selection in subgenome-B versus nine in subgenome-A (**Fig. 5b**). While there is no strong evidence of rediploidization or marked subgenome dominance, genomic redundancy has allowed slightly deleterious and beneficial mutations to accumulate. Whether redundancy will foster adaptation and long-term resilience in this homostylous, narrow endemic species remains an open question.

## MATERIALS AND METHODS

### Genome-size estimation, DNA and RNA isolation and sequencing

Genome size was estimated via flow cytometry in three *P. grandis* individuals according to Potente et al. (2022a). Additionally, we estimated the genome size using *k*-mer frequency spectra from raw long-read sequencing (see below) with GenomeScope 2.0 (Ranallo-Benavidez *et al*., 2020). To generate sequencing libraries, DNA was extracted from flash-frozen leaves of one individual following a modified CTAB protocol (Potente *et al*., 2022a). Paired End (PE) 150bp reads were generated via Whole Genome Sequencing (WGS) with Illumina (∼45x coverage) at Functional Genomics Center Zurich (FGCZ, Switzerland). Additionally, long-read sequencing was performed on one PromethION flow cell at FGCZ and three MinION f Mk1B flow cells in-house, achieving a combined ∼35x coverage. Long reads were basecalled with Guppy v.6.0.1 specifying the Super accurate (SUP) model. A Hi-C library was prepared following the protocol by Grob et al. (Grob *et al*., 2014), then sequenced on Illumina NovaSeq 6000 (PE 150bp reads; ∼375x coverage) at FGCZ.

Transcriptomes from leaves, flowers and floral buds of a single individual were obtained using the Spectrum Plant Total RNA Kit (Sigma-Aldrich). Subsequently, TruSeq Stranded mRNA libraries were prepared and sequenced on NovaSeq 6000 at FGCZ, generating ∼120 million PE 150bp reads per sample.

### Genome assembly and annotation

Genome assembly was performed with MaSuRCa v.4.0.5 (Zimin *et al*., 2017); Hi-C reads were mapped to the resulting reference genome using BWA (Li & Durbin, 2009) and scaffolded with YaHS v.1.2a.1 (Zhou *et al*., 2023); Hi-C contact map visualization and manual curation of mis-assemblies were achieved with JuiceBox (Durand *et al*., 2016). *Primula grandis* chromosome-sized scaffolds were ordered and named based on collinearity with *P. veris* scaffolds.

A set of protein sequences for homology-based annotation was prepared by merging the OrthoDB protein data set for Viridiplantae (odb10) with a high-quality *P. veris* protein set that excluded monoexonic genes as performed by Potente et al. (2024). Then, Braker3 was used to annotate the genome using the RNA-seq alignment file. We ran BUSCO v4.0.6 (Manni *et al*., 2021) on the coding sequences of *P. grandis* using the eudicot database (odb10) to assess gene-annotation completeness. Identification and annotation of repetitive elements was performed using Extensive De novo TE Annotator (EDTA) v1.9.2 (Ou *et al*., 2019), as in Potente et al. (2024) (SI Appendix).

### Subgenome characterization

If *P. grandis* is allotetraploid, as proposed by previous phylogenetic analyses (Schmidt-Lebuhn *et al*., 2012; Stubbs *et al*., 2023), it should contain two distinct subgenomes. We tested this prediction by using subgenome-specific *k*-mers-analyses of LTRs implemented in SubPhaser v.1.2.6 (Jia *et al*., 2022). To test for subgenome dominance, we checked for Homoeologous Expression Bias (HEB) towards one of the subgenomes of *P. grandis* as follows. First, we aligned transcriptomic data from leaves, flowers and floral buds to the reference genome using Bowtie2 v2.3.4.3 (Langmead & Salzberg, 2012). Secondly, the number of aligned reads mapping to each gene was counted using htseq-count v2.0.5 (Putri *et al*., 2022) to estimate Reads Per Kilobase per Million (RPKM). Thirdly, HEB was estimated as the log_2_ difference between RPKM of homoeologs from subgenome-A and -B, respectively, using −3 ≤ log_2_ HEB ≥ 3 (i.e., 8-fold HEB bias towards either subgenome-A or -B) as significance threshold (Smith *et al*., 2019). Finally, we estimated the divergence times between *P. grandis* subgenomes and between each *P. grandis* subgenome and *P. veris* genome via the estimation of genome-wide divergence at synonymous sites (*Ks*) (see SI appendix). To infer absolute ages for genome divergence among subgenomes, we used the neutral substitution rate of 6.15 × 10^−9^ substitutions per synonymous site per year previously determined for *P. veris* (Potente *et al*., 2022a).

### *S*-locus identification and tests of selection on *S*-genes

To discover whether the *S*-locus is found in only one or both subgenomes of *P. grandis* and identify its genomic location, structure and organization, we performed microsynteny analyses of *P. veris S*-locus against the *P. grandis* reference genome using MCscan (Tang *et al*., 2008). To determine the number of *S*-locus copies in the *P. grandis* genome, we estimated sequencing coverage analysis of the *S*-locus and its up- and down-stream regions using WGS data from 10 *P. grandis* individuals (see below). To assess whether the loss of heterostyly in *P. grandis* is associated with *S*-genes degeneration, their sequences were extracted from the reference genome using bedtools v2.29.2 (Quinlan & Hall, 2010) and aligned to the coding sequences of functional *P. veris S*-genes using the MUSCLE alignment algorithm as implemented in MEGA X (Kumar *et al*., 2018). Moreover, to determine the levels of *S*-locus genetic diversity, we estimated nucleotide diversity (π) in all *S*-genes (SI Appendix) of ten *P. grandis* individuals (see below). Finally, to ascertain whether selection on *P. grandis S*-genes is relaxed, we performed McDonald-Kreitman (MK) tests by comparing the S-genes of ten *P. grandis* individuals with those of six *P. veris* individuals (see below; SI appendix).

### Comparative genome-wide analyses

To perform comparative genomic analyses, we used a WGS data set comprising 10 newly sequenced individuals of the Caucasian endemic *P. grandis* and 80 previously sequenced individuals from the Caucasus of the six diploid species in *Primula* sect. *Primula* (*P. veris* subsp*. macrocalyx*, *P. vulgaris* subsp*. vulgaris, P. elatior* subsp*. pallasi*, *P. megaseifolia*, *P. renifolia*; (Stubbs *et al*., 2023); Table S1; SI Appendix). First, reads were trimmed with Trimmomatic v0.38 (Bolger *et al*., 2014). Secondly, the reads of each individual were mapped once to each subgenome of *P. grandis* separately using BWA-mem v7.17 (Li & Durbin, 2009). This latter decision was taken due to the fact that the above species are diploid, whereas the newly assembled genome is tetraploid. Therefore all subsequent analyses were performed separately for each subgenome. Then, variant calling was performed using the functions *mpileup* and *call* included in BCFtools v1.8 (Danecek *et al*., 2021). Variant sites were filtered with VCFtools v0.1.14 (Danecek *et al*., 2011) and BCFtools.

*Demographic analyses* - To test the expectation that polyploid speciation is associated with a genetic bottleneck, we inferred single-population demographic histories of *P. grandis* using Stairway Plot v2 (Liu & Fu, 2020). Furthermore, to estimate the timing of the genetic bottleneck expected with *P. grandis* speciation, we performed model-based coalescent simulations of demographic changes with fastmsimcoal2 v2.7 (Excoffier *et al*., 2021) using 2D-Site Frequency Spectrum (2D-SFS; see SI Appendix). Finally, we compared the demographic history of *P. grandis* subgenomes-A and -B with its relatives in *Primula* sect. *Primula* by estimating genome-wide distributions of Tajima’s D in all species.

*Phylogenetic analyses* - To determine whether any extant species in *Primula* sect. *Primula* served as progenitors of *P. grandis,* we first identified collinear orthologs shared between both subgenomes of *P. grandis* and the assembled genome of *P. veris* using MCscan (github.com/tanghaibao/jcvi/wiki/Mcscan-(Python-version); (Tang *et al*., 2008)). Using the concatenated ortholog dataset, we then inferred the Maximum Likelihood (ML) phylogeny of *Primula* sect. *Primula* using IQ-TREE v.2.1.2 (Nguyen *et al*., 2015) and estimated statistical support for each node with 1000 ultrafast bootstraps (BS) (Minh *et al*., 2013; Chernomor *et al*., 2016).

*Population genomic analyses and genome-wide tests of selection*- To test the expectations of genome-wide reduced genetic diversity and increased Linkage Disequilibrium (LD) associated with polyploid-speciation and shift to self-compatibility, we compared the number of segregating sites (S), and nucleotide diversity (π) at both synonymous and non-synonymous sites using the python script ‘popgenWindows.py’ (https://github.com/simonhmartin/genomics_general) and LD decay using popLDdecay (Zhang *et al.,* 2019) across the genomes of the tetraploid, homostylous *P. grandis* vs. its diploid, heterostylous relatives in *Primula* sect. *Primula* (**Fig. 5a**; see SI appendix). To test the hypothesis of genome-wide relaxed selection due to increased genetic redundancy in polyploid genomes, we compared the strength and Direction of Selection (DoS) in the genes of each *P. grandis* subgenome vs. its diploid relatives using the McDonald-Kreitman (MK) tests as implemented in degenotate.py (https://github.com/harvardinformatics/degenotate; SI appendix).

## DATA AVAILABILITY

Raw sequencing data (long-read and short-read sequencing and Hi-C data) for the genome assembly of *P. grandis* will be uploaded to the NCBI SRA upon acceptance. at The 90 whole genome sequences used in this study are deposited in SRA (BioProject ID: PRJNA1066534). Code and R scripts used to run the programs and create figures will be available in (https://github.com/EmilianoMora/primula_grandis_project).

## ACKNOWLEDGEMENTS

We thank the Functional Genomics Center UZH for help during library preparation. This research was supported by the Swiss National Science Foundation grant nos. 3100-061674.00/1 and 31003A_175556 to EC.

